# The limitations of *in vitro* experimentation in understanding biofilms and chronic infection

**DOI:** 10.1101/032987

**Authors:** Aled E. L. Roberts, Kasper N. Kragh, Thomas Bjarnsholt, Stephen P. Diggle

## Abstract

We have become increasingly aware that during infection, pathogenic bacteria often grow in multicellular biofilms which are often highly resistant to antibacterial strategies. In order to understand how biofilms form and contribute to infection, *in vitro* biofilm systems such as microtitre plate assays and flow cells, have been heavily used by many research groups around the world. Whilst these methods have greatly increased our understanding of the biology of biofilms, it is becoming increasingly apparent that many of our *in vitro* methods do not accurately represent *in vivo* conditions. Here we present a systematic review of the most widely used *in vitro* biofilm systems, and we discuss why they are not always representative of the *in vivo* biofilms found in chronic infections. We present examples of methods that will help us to bridge the gap between *in vitro* and *in vivo* biofilm work, so that our bench-side data can ultimately be used to improve bedside treatment.

## Introduction

Bacteria were once thought to exist as single, free-floating planktonic cells that are community independent. John William (Bill) Costerton changed this perception in the late 1970s when he observed surface associated microbial aggregates enclosed within a matrix of extracellular material, a phenomena he later termed ‘biofilm’ [1,2]. Today, the biofilm phenotype has been identified in up to 80% of all non-acute infections, including foreign-body related, otitis media, orthopaedic, catheter, chronic wounds and lung-related infections [3–7]. The interchange between planktonic and biofilm phenotypes, is believed, but not proven, to commonly manifest clinically as acute and chronic infections respectively.

Acute infections tend to be fast spreading with a rapid onset. They are often controlled by the host immune response, and excessive intervention is not always required. However, if the host defences fail and therapeutic intervention is required, acute infections can usually be cleared within days [8]. Conversely, chronic infections are where there is a delay in the healing process (an inability of the injured site to restore anatomical and functional integrity), consistent with the severity of the injury [9]. The presence of biofilms and their innate ability to tolerate antibiotics up to 1000 times greater than planktonic cells, is thought to delay wound restoration [10–13]. Cells assuming the biofilm phenotype are commonly observed in patients with various underlying conditions, which can be system wide in the case of immunodeficiency and diabetes, or more focused in the case of venous leg ulcers and cystic fibrosis (CF). In the case of CF, chronic biofilm infections have been known to persist in the airways for over 30 years. Therefore, chronic infections are an ever increasing problem due to their recalcitrance towards extensive antibiotic treatment regimes and persistence under sustained attack from the host’s innate and adaptive immune response systems [10,14].

The treatment and management of patients suffering from chronic infections represents a significant monetary and labour intensive burden to healthcare providers. Recently, the direct costs of chronic infections, such as those affecting the dermis, were estimated to be in excess of $18 billion, affecting 2 million residents in the United States alone, and resulting in 200,000 deaths annually [15]. This data relates to one country, and a single infected organ. If this is representative of other countries, and other conditions, then chronic infections represent a huge worldwide problem. A recent report on antimicrobial resistance (AMR), has stated that we can expect up to 10 million extra deaths annually worldwide by 2050 due to AMR [16]. The problem of chronic infection is only going to aid the rise of AMR.

Much of our current knowledge about chronic infection comes from studying bacteria growing in test tubes. Costerton encouraged laboratories worldwide to deviate from studying planktonic cultures, and instead, focus on understanding surface associated biofilms. This has become increasingly more relevant as we battle with the issues posed by AMR. We are becoming increasingly aware of significant differences that exist between *in vitro* biofilms grown in the laboratory, and *in vivo* biofilms found during actual infection. This raises the question as to whether the experiments that we currently perform in the laboratory are useful for understanding how bacterial biofilms form and contribute to AMR during infection.

To understand the biology of infection better, we need another paradigm shift, a new wave of methods and experiments that better represent clinical conditions (of which biofilm formation is only one aspect). The use of some methods can potentially hamper our understanding of various aspects of infection, as they do not always accurately represent what we observe clinically. This review summarises what we know about bacteria during infection, and how our current *in vitro* methods fail to represent such factors. In the following sections we discuss biofilms and polymicrobial interactions, particularly in the context of *Pseudomonas aeruginosa* as one of the most common opportunistic pathogens that causes chronic infection. We explore the differences between *in vitro* and *in vivo* observations, and discuss how to better bridge the gap between the two, increasing experimental accuracy so that our bench-side data can be used to improve bedside treatment.

## The role of biofilms during chronic infection

Chronic infections persist despite apparently adequate antibiotic therapy, and in the face of the host’s innate and adaptive defence mechanisms. Chronic infections are characterised by persistent and progressive pathology, mainly due to the inflammatory response surrounding *in vivo* biofilms [17]. This biofilm lifestyle appears to impair the host’s ability to combat the infectious agent. The innate immune response in the form of recruitment of neutrophils and their inability to break through the biofilms defence has been specifically examined [18–21]. Polymorphonuclear leukocytes (PMNs) are recruited in large numbers to the infection site, and during acute infection, are able to phagocytize and remove most of the infectious agent [3,18]. When PMNs fail to eradicate an infection, it is most likely because a biofilm has been established. In the biofilm state, the enclosing matrix of extracellular substances is capable of protecting underlying cells from the immune system, such as PMN phagocytosis [22,23]. In addition to this, biofilms are capable of suppressing the antimicrobial action of PMNs through the production of various virulence factors [19,24]. *P. aeruginosa* growing in biofilms has been shown to actively kill PMNs through the secretion of rhamnolipids which reduce the host’s ability to clear infection [25].

In contrast to acute infections, which are usually treatable by traditional antibiotics, biofilms are known to tolerate antibiotic concentrations up to 1000 times higher than the minimal inhibitory concentrations (MIC). It is important to note that there are differences between antibiotic resistance and tolerance, with many studies reporting the former when meaning the latter. Antibiotic resistance is where there is an increase in the MIC through mechanistic intervention. Conversely, little is known about antibiotic tolerance. There is often little change in the MIC of individual cells, however growth in populations can lead to an increasing tolerance to antibiotics. The cause of such increased tolerances seen in biofilms has been investigated in a number of different studies, focusing on the extracellular matrix, the involvement of quorum sensing (QS), and the physiological factors observed within the biofilm [19,26–29]. The extracellular matrix itself has been shown to obstruct the diffusion of some antimicrobial agents through both chemical and physiological means [27,30,31]. The matrix however is not selective against antimicrobial agents, and therefore the diffusion of many substances, such as oxygen, metabolites and waste products, can also be altered. Reduced diffusion in combination with high cell densities results in steep chemical gradients from the outer surface towards the central core of an *in vitro* biofilm [26,32]. This causes a very heterogenic growth pattern throughout *in vitro* biofilms, which in turn influences regional tolerance towards different types of antibiotic treatments [33,34]. Tobramycin tolerance in *P. aeruginosa* has also been shown to be influenced by QS-systems [19]. These studies have predominantly used *in vitro* systems in an attempt to explain the increased persistence and tolerance of biofilm cells towards varying antibiotic treatment regimens, and the failures of the host immune system to eradicate infection. However, such studies are very difficult to transfer directly to *in vivo* settings.

Another observation of chronic infection, is the presence of multiple species within an infection site [35,36]. The milieu of species within a defined space may result in cooperation and/or conflict with other community members [37–39]. This creates a complex environment where species align along different nutrient gradients, be those host derived or the result of species interactions, as seen between *P. aeruginosa* and *Staphylococcus aureus* in chronic wounds [40–42]. The complexity of interactions between species may be enhanced through the production of diverse QS signals within the infection site, which have inter- and intra-species effects, although this is yet to be demonstrated *in vivo* [43–45]. One thing of note, is the lack of evidence supporting multi-species biofilms during infections, whereby species grow concomitantly within the same biofilm, however the presence of multiple species within an infection site is well documented.

## *In vitro* investigation of biofilms

During the last three decades, biofilms of pathogenic species have been extensively studied by a wide range of research groups, each with differing objectives, but all with the same overall aim – to expand our knowledge of biofilms to better understand infection. Using continuous flow-cell conditions and Confocal Laser Scanning Microscopy (CLSM), we are closer to understanding the processes involved in the initial attachment of cells to surfaces *in vitro* [46–48]. Combining molecular techniques with CLSM to construct and visualise knock-out strains of bacteria, has shown the contribution of motility and QS to biofilm development [30,33,46,47,49].

The development of low cost, high throughput biofilm screening methods in microtitre-plates, have made it possible to identify genes essential for surface-attached biomass production in liquid media [50,51]. This system has been heavily used to identify potential anti-biofilm agents, by measuring the reduction in surface attached biomass on the sides of wells after treatment with potential therapeutic agents [51–54]. In addition to this, The Center for Disease Control (CDC) approved biofilm reactor [55–57], and drip flow reactors, have proved excellent for assessing biofilm formation on biological and non-biological materials [58–60]. These flow systems and microtitre assays are the workhorses of *in vitro* biofilm research, and have generated a phenomenal amount of data that has greatly expanded our knowledge about how bacteria attach and differentiate into mature biofilms *in vitro*.

The commonality between these methods is the growth of biofilms on abiotic surfaces, that are submerged in media, and exposed to fluid dynamics of varying degrees. Experiments in these systems are able to produce biofilms of high cell density with a topography that can reach up to several hundred μm thick. Under continuous flow cell conditions, the emergence of mushroom structures and water channels occur after only a couple of day’s growth (Figure 1) [46,47]. A key question is whether we are able to transfer our *in vitro* knowledge from the laboratory bench to the patient bedside, and the last decade has provided us with refined techniques that allow us to gain insight into what is actually occurring within certain types of infection.

**Figure 1.**
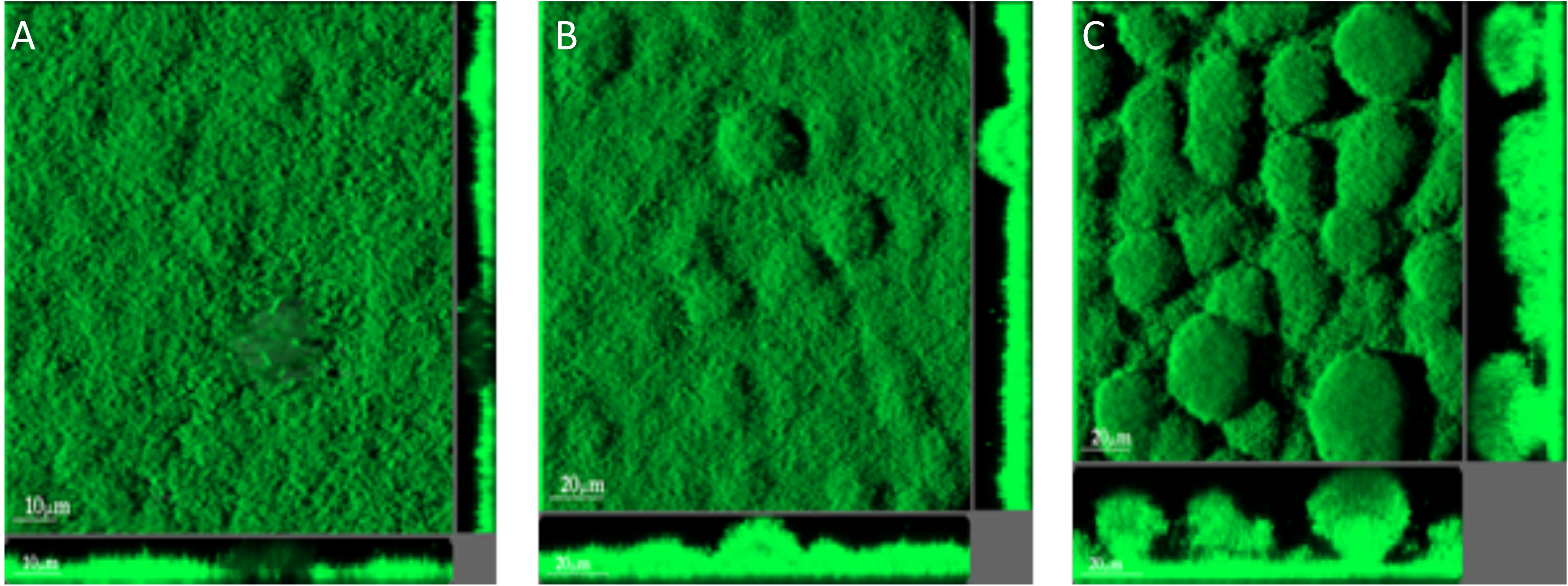
Confocal laser scanning micrographs of 2-day-old (A), 3-day-old (B) and 4-day-old (C) biofilms formed by a *P. aeruginosa* wild-type strain in a continuous flow cell system. The central images show top-down views, and the flanking images show vertical optical sections. The bars represent 10 μm (A) and 20 μm (B and C). Bjarnsholt, unpublished.

With advanced microscopy techniques, we are now able to see how bacterial cells organise themselves in chronic infections in *ex vivo* samples. As with the majority of biofilm research, there has been considerable focus on infection in patients with CF. In *ex vivo* lung tissue infected with the major CF pathogen *P. aeruginosa*, much of the bacterial biomass is found as a biofilm within the bronchial lumen. Interestingly, these biofilms have not been found attached to the epithelial surface, but are found as non-attached biofilm aggregates that are embedded in the highly inflamed mucus [3,21] (Figure 2). Lying completely surrounded by immune cells, mainly PMNs, these aggregates range in size from 4 μm – 100 μm in diameter [3,20]. The same mode of non-attached aggregated biofilm growth have been observed in other types of infection, such as those found in chronic wounds, otitis media, and soft tissue fillers [4,5,61].

**Figure 2.**
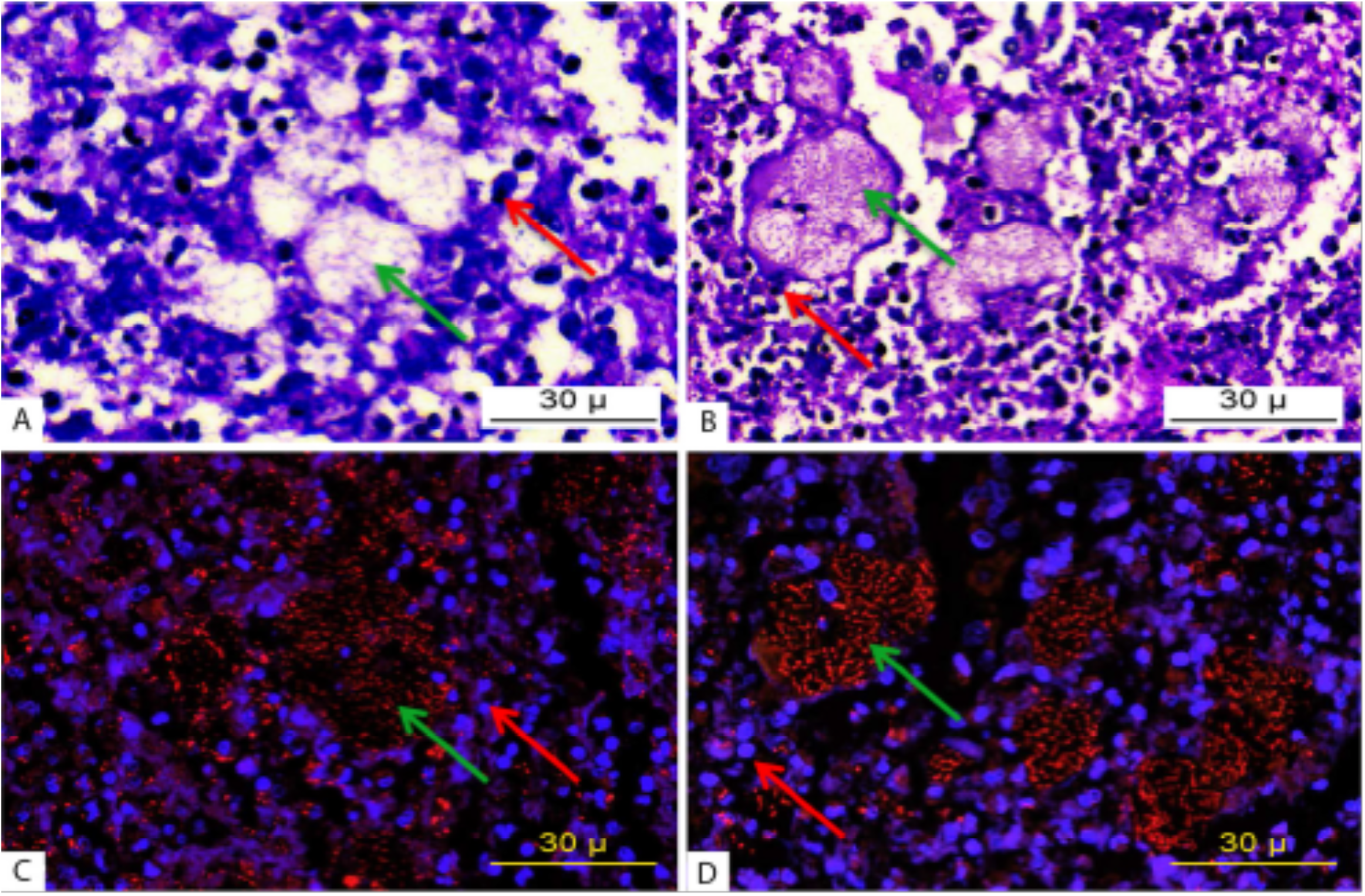
Micrograph of *P. aeruginosa* infected lung tissue from a patient with cystic fibrosis. Light and fluorescence microscopy images (170× magnification) of PAS hematoxylin-stained (A, B) and PNA FISH-stained (C, D) sections containing luminal and mucosal accumulations of inflammatory cells. The *P. aeruginosa-positive* areas are seen as well-defined lobulated clarifications surrounded by inflammatory cells. Red arrows indicate PMNs and green arrows indicate *P. aeruginosa* biofilm aggregates [21].

A glaring discrepancy between *ex vivo* observations and the *in vitro* biofilm, is the absence of a surface. In all of the infections mentioned above, attachment to a surface or the epithelia is rarely observed. However, the majority of primary *in vitro* models for biofilm infection have surface attachment as a crucial component. Recent studies have shown how *P. aeruginosa* non-attached biofilm aggregates form in liquid batch cultures, and that they have the same characteristics as surface attached biofilms when it comes to antibiotic tolerance and resilience towards PMNs [62,63]. The absence of such surfaces in surface wounds means the bacterial cells assume a biofilm lifestyle within a self-contained aggregate, devoid of surface attachment [64,65]. There have been a number of *in vitro* studies that have observed steep chemical gradients (notably oxygen) throughout biofilms, which leads to a heterogenic growth pattern [26,32]. The complex environment inside inflamed mucus seems to govern the growth pattern in biofilms in a different way. When the growth rate of aggregated bacteria within mucus is measured, a heterogenic growth pattern is observed, due to large regional variations between aggregates which can be linked to the local concentration of PMNs [21]. Measurements of expectorated sputum from CF patients with a *P. aeruginosa* infection, demonstrates a micro-aerophilic environment as a result of oxygen consumption by PMNs [66]. These findings point towards immune cells playing a central role in regulating the growth of bacterial cells as they grow in aggregates, at least during infection in the CF lung. Crucially, immune cells are absent in almost all *in vitro* systems.

Some prominent biofilm infections do involve the “characteristic” attachment and growth of biofilm cells on a surface. Chronic foreign body infections of implants, stents and catheters, result in bacterial cells colonizing abiotic surfaces, leading to biofilm formation [67–69]. Catheters have been shown to constitute an environment whereby bacterial infections are able to develop into very thick biofilms [67,68,70]. Therefore, catheter associated biofilms are an example of a biofilm that most resembles the flow of media over a surface, and are most similar to CDC reactors, drip flow and flow cell systems. There are still some striking differences. For example, the type of structures observed and the level of organization seen in flow cells, are not present in catheter related biofilms, and mushrooms structures are almost never observed. Other differences include the liquid flowing over the surface. During *in vitro* biofilm growth, well-defined minimal medias are most often used [49,56,57,71,72]. In catheters, the liquids flowing over the surface could be urine, blood or other bodily fluids, which constitutes a very different growth environment for bacteria. In many other biofilm infections, there is a distinctive lack of flow within the infection site [20]. However, the lack of a flow does not restrict the bacteria from creating a biofilm. For example, *P. aeruginosa* is capable of biofilm formation deep within the dehydrated mucus of the CF lung, or in the inflamed pus of chronic wounds [73,74].

The pros and cons of each of these methods are listed in Table 1. Many of these methods produce clear and well defined results, due in part to our ability to control experimental parameters with a high degree of stringency, whilst concomitantly allowing single variables to change. This allows us to study the effects of single elements on various aspects of biofilm growth. This reductionist approach has yielded great information about complex cell matters such as metabolism, resistance mechanisms, and signalling pathways. However, such a simplistic approach is not always possible during *in vivo* methodologies, due to natural variations between living organisms.

**Table 1.** Comparison of different model systems for studying biofilms *in vitro*.

| Method | Description | Pro's | Con's | Ability to reflect chronic infections | References |
| --- | --- | --- | --- | --- | --- |
| **Microtitre assay/**<br>**Calgary model** | Test the buildup of biomass on pegs or the sides of wells filled with static media | • Very high throughput screening<br>• Cheap and simple methodology | • Does not differentiate between matrix, living and dead cells attached to the surface<br>• Large biological variations between wells | • Does not reflect any environment observed in the human body<br>• Surface attachment is only found in foreign body infections<br>• Does not reflect complex oxygen and nutrient gradients found in infections<br>• Missing immune response | [50,51] |
| **CDC Reactor** | Test biofilm growth on disks spinning around in a chemotactic media | • Able to test biofilm growth on diverse types of material<br>• Tablets can be removed for microscopy and CFU counting<br>• High reproducibility | • High shear forces on the disk may remove biomass | • Surface attachment is only found in foreign body infections<br>• Does not reflect complex oxygen and nutrients gradients found in infections<br>• Missing immune response. | [56,59] |
| **Continuous Flow Cell System** | Growth of biofilm on glass supplied with a continuous flow of low carbon content media | • Enables structural analysis of biofilm growth by CLSM<br>• Biomass can be quantified by microscopy<br>• Produces 20-100 $\mu\text{m}$ thick highly structured biofilms | • Complex & expensive system which requires a CLSM<br>• Media most often used are well defined minimal medias<br>• Low reproducibility and high variations within same chamber<br>• Labor and time consuming | • May reflect some infections such as urinal infections<br>• The structural 'mushroom' biofilm seen in flow cells have never been observed during infection<br>• Does not reflect complex oxygen and nutrients gradients found in infections<br>• Missing immune response | [72,143] |
| **Drip Flow Reactor** | Drops of media continuously feeding the growth of biofilms on an object glass placed at an angle | • Very high yield of biofilm<br>• May be used both for direct microscopy, CFU counting or transcriptomic analysis<br>• Produces several 100 $\mu\text{m}$ thick biofilms | • High shear forces in the drip area of impact<br>• Very messy and varying biofilm<br>• Uses large volumes of media | • Surface attachment is only found in foreign body infections<br>• Does not reflect complex oxygen and nutrients gradients found in infections<br>• Missing immune response. | [57,58] |

## *In vivo* investigation of biofilms

More recently, the advent of *in vivo* methods have increased our understanding of biofilms during chronic infection. There are a range of *in vivo* models that simulate chronic infections, such as surface wounds [75,76], subcutaneous wounds [77,78], implant-related (such as catheter, orthopaedic, and dental) [79–83], otitis media [84–86], and CF [87–90] to name but a few. As with all models, some are deemed more applicable than others. For instance, porcine models lend themselves greatly to the study of chronic wound infections due to similarities of the immune response systems, spatial structuring within tissue, and wound healing processes (re-epithelialisation, scarring, and tissue granulation) [91,92]. The complexities within *in vitro* biofilms such as structure, gene regulation, and the production of virulence factors, has been elucidated for many problematic opportunistic pathogens. However, during *in vivo* chronic infection, there is a complex interplay between host and pathogen, which is not easily replicated *in vitro*, and leads to observable differences between *in vitro* and *in vivo* “chronic infections”.

The *in vivo* biofilm differs from its *in vitro* counterpart, in both size and “shape”. A meta-analysis on the size of *in vivo* biofilms from chronic infections by Bjarnsholt and colleagues (2013) [20], showed them to have an upper size limit of 200 μm, which can be superseded in the presence of abiotic surfaces (e.g. catheters) leading to biofilms in excess of 1000 μm. Such observations differ drastically to the swathes of biofilm growth (up to several cm^2^) observed using *in vitro* methods [49]. It is thought that size limits placed on *in vivo* biofilms are the result of limiting factors, once thought to be nutrient based, but evidence points to oxygen depletion in the local environment [93]. In addition, biofilm “shape”, or more accurately the 3D structure, is different under *in vitro* conditions, with the characteristic mushroom structures of *P. aeruginosa* biofilm formation, yet to be observed *in vivo*.

Following periods of trauma, the natural microbial flora may develop into antagonistic biofilms and a state of chronicity. *In vivo* models of chronic infection require artificial inoculation, usually at an inflated concentration, and with the aid of foreign bodies, so that the inoculation is not cleared by the host immune system [82,94,95].

Another difference between many *in vivo* studies and actual *in vivo* infections, is the potential for multiple species to be present within the latter. A meta-analysis of 454 wound biofilms by Peters and colleagues from diabetic patients identified more than 1600 unique bacterial species, with diversity similar to that of the patients natural skin flora [35,96]. The presence of two or more unique bacterial species is observed in more than 80% of wounds analysed, with a relatively large proportion (30%) containing five or more species. Other studies place diversity higher, with an average of 5.4 bacterial species per chronic wound [36]. Such degrees of species diversity are also observed in the CF lung, resulting from the diverse microenvironments that arise [97–100]. Conversely, Kirketerp-Moller and colleagues (2008) [5] struggled to identify multiple species during a venous leg ulcer study, highlighting the diversity and complexity of different types of chronic infection. For instance, *P. aeruginosa* is known to grow in the deeper recesses of chronic leg ulcers, compared to *S. aureus*, which is found on the surface [42]. This non-random distribution suggests some species, especially anaerobes in deep recesses, will be missed if the wrong sampling method is employed. Therefore, an all-encompassing, biopsy-based extraction and molecular identification, should be employed to allow a thorough investigation of species within an infection site.

This degree of diversity within an infection site may result in a range of species-species interactions that have the potential to be both beneficial and antagonistic. For instance, *Haemophilus influenzae* and *Moraxella catarrhalis* grown concomitantly in an otitis media Chinchilla model, show an increased tolerance towards antimicrobial agents and the host’s response system [101]. In other models, differences are also observed. For instance, co-colonisation of two common CF pathogens (*P. aeruginosa* and *Burkholderia cenocepacia*) in a murine CF model leads to a mutualistic relationship, whereby *P. aeruginosa* persists, and *B. cenocepacia* alters the inflammatory response [102]. In some instances, the presence of multiple species within a specified environment is a necessity. For instance, during colonization of the oral cavity, primary colonisers (such as *Candida albicans*) allow the attachment of other species (such as *Streptococcus spp*.), facilitating the temporal co-aggregation of cells [103]. Should one of the primary colonisers be absent, co-aggregation will cease. Whilst highlighting the need to study multi-species infections, it also suggests that we need to incorporate a diverse range of species into experiments. It is important to re-iterate that the presence of multiple species within a system (such as an infection or test tube) does not mean they form a multi-species biofilm, which has not been observed to date. Therefore, it may be best to think about such systems as “battlefields”, with species not directly mixing, but residing within their own ecological space.

Whilst some species act synergistically with others through metabolite and signal production, and/or direct contact, some species have been observed to diversify without the need for multi-species interactions. For instance, during CF lung infection, *P. aeruginosa* gradually evolves, adapting to the CF lung through structural and dynamic changes over time that reduce virulence, and increase chronicity through a phenotypically heterogeneous population [104]. Interestingly, seemingly homogenous subpopulations of *P. aeruginosa* in the CF lung show large variation for many key aspects of infection, such as antibiotic tolerance, QS and virulence factor production [105–108]. Such diversity between and within species, may alter the pathogenicity of the infection, infection persistence, impact on the host and antibiotic resistance and tolerance.

To better understand the effects of biofilm infections on wound healing, both murine and rodent models have been devised [109,110], however there remains a disparity in their effectiveness due to the differences seen during wound healing in humans. As mentioned previously, porcine models can provide increased translational data for delayed wound healing in light of biofilm infection due to similar dermal properties (re-epithelialization, scarring, hair follicle placement and abundance), however their accessibility is more restrictive and expensive than that of murine/rodent models [111]. Implant models are a great way to blur the lines between *in vitro* and *in vivo* biofilms through the presence of an abiotic surface. Comparing *in vivo* models where an abiotic surface is present to *in vitro* biofilms, results in very similar biofilms (size, shape, and thickness) [20]. There are a range of *in vivo* models for chronic lung infection, however most require the require the bacterial cells to be well established on agar/agarose sheets or attached to the surface of alginate beads [95]. A murine model whereby cells are inhaled, has been developed to circumvent the need for various inoculation implants, whilst concomitantly allowing the upper and lower respiratory tract to be investigated [112]. Further to this, and more importantly, the model allows for persistence and adaptation of cells, similar to that observed in chronic infections, which includes evolutionary dynamics. The range of *in vivo* methods, along with their pros and cons have been summarised in Table 2.

**Table 2.** Comparison of different model systems for studying biofilms *in vivo*.

| Method | Description | Pro's | Con's | Ability to reflect Chronic infections | References |
| --- | --- | --- | --- | --- | --- |
| ***Porcine models*** | Especially used to simulate chronic wounds | • Similar immune response to skin infections<br>• Similar skin structure<br>• Large skin surface | • Very expensive<br>• Space consuming<br>• Opportunistic pathogens may vary<br>• Ethical considerations | • Closely related to human wound healing<br>• Immune response towards very similar to human<br>• Are able to reflect complex oxygen and nutrients gradients found in infections | [91] |
| ***Murine models*** | Have been used for chronic wounds, CF and foreign body infections | • Less expensive than porcine<br>• Small space needs<br>• Fast generation time | • Very hard to maintain chronic infections<br>• High metabolism<br>• Ethical considerations | • Murine models have a very different immune response compared to humans<br>Most infections clear fast.<br>• Needs modified environment (implant, inserted beads etc.) or a reservoir (via natural inhalation) to facilitate prolonged infection<br>• Are able to reflect complex oxygen and nutrients gradients found in infections | [79,82,86,109,112] |
| ***Other rodents*** | Rats, rabbits, chinchillas etc. have been used as otitis media, catheter, osteomyelitis models | • All of these models can be excellent to examine infections in complex environments<br>• Specialized models (chinchilla otitis media model or Rabbit burn wound model) | • Expensive and labor intensive<br>• Ethical considerations<br>• Very considerable reservations as to how closely they resemble human infection | • Rodents generally have much higher metabolism and pulse compared to humans<br>• Rodent models have a very different immune response compared to humans<br>• Implant infections show much the same biofilm buildup as observed on infected human implants<br>• Are able to reflect complex oxygen and nutrients gradients found in infections | [81,144,145] |

## *In vivo* conditions, *in vitro* methods

Sometimes it may not be ethical, practicable, or feasible to conduct *in vivo* experimentation. Given the issues highlighted previously, how can we better represent *in vivo* conditions in our *in vitro* models? It is widely known, for *P*. *aeruginosa* at least, that different nutritional cues result in altered biofilm formation, virulence, motility, and QS [46,113–117]. These differences become increasingly important when factors of clinical relevance, such as virulence and antimicrobial tolerance, are altered [13,118,119]. Nutritional cues similar to those of expectorated CF sputum have been incorporated into a synthetic CF sputum media (SCFM) that approximates *P. aeruginosa* gene expression to that observed in expectorated CF sputum [120]. Two noteworthy points of this study are (i) the lack of key components observed in CF sputum (notably DNA, fatty acids, N-acetyle glucosamine, and mucin) and (ii) the inability of the methods used (RNA-seq) to correctly predict fitness requirements [121–127]. A follow up study rectified these issues, incorporating these components into SCFM. Using Tn-seq (which is a more precise way to measure fitness, compared to RNA-seq), it was shown that near identical selection pressures exist between synthetic and expectorated CF sputum [128]. In a separate study, the use of an artificial sputum media was shown to increase diversity within a population, something not observed in Lysogeny Broth (LB) [108]. This diversity increased in the presence of certain antibiotics at sub-inhibitory concentrations, highlighting the need for effective clearing of cells. Whilst the use of specialized media will not replace *in vivo* techniques, any way in which we can manipulate cells to generate increased diversity as seen during *in vivo* experiments, may result in findings that have increased clinical relevance.

Similarities have been observed in the active biosynthetic pathways of *P*. *aeruginosa* in the CF lung and murine surgical wound infections [127]. This suggests that (i) catabolite metabolism is shared between certain infection sites and (ii) other factors, such as host-inflammatory responses, may be the cause of infection chronicity. In light of this, it might be better to think of SCFM, not as a synthetic sputum media, but a synthetic infection media (SIM), which could then be supplemented further to better replicate the nutrient environment of *in vivo* conditions. For example, surface wounds contain high concentrations of both host-derived serum proteins and the fibrous extracellular matrix protein; collagen [129,130]. The presence of such compounds in growth media is known to reduce biofilm formation for a range of clinically relevant organisms, including *P. aeruginosa* and *S. aureus* [131–133]. The problem with such observations is that the experiments were performed using the microtitre assay, which does not take into account the possibility that attachment (to abiotic surfaces) might be the cause of such observations, and which under many *in vivo* conditions has little relevance. Such observations may also suggest why surface attachment (to biotic surfaces) is not generally observed *in vivo*, and why complex 3D structures do not develop.

*In vitro* techniques are widely criticized for their incorporation of abiotic surfaces, which only have clinical relevance to a small number of implant-related infections [67–69]. However, the widespread use of abiotic surfaces is not surprising due to the difficulties of trying to mimic the complex multicellular topology of an *in vivo* surface. The easiest way to negate these complexities is to employ surface independent methodologies, which have been shown to produce biofilms of equal size, shape, and antimicrobial tolerances to those observed *in vivo* [62,133,134]. Whilst these methods produce *in vivo* biofilms under *in vitro* conditions, they lack the complex 3D topology and spatial structure of host tissue, which will alter a range of factors from antimicrobial to nutrient and oxygen penetration [21,135,136].

Negative effects asserted on biofilm formation by serum proteins are also observed for antibiotic penetration in tissue samples [136]. As mentioned previously, bacteria are capable of occupying different ecological niches within a wound environment [42]. Growth of different methicillin-resistant *Staphylococcus aureus* (MRSA) strains on porcine nasal epithelium tissue resulted in three different growth profiles, highlighting the need to select strains with clinical relevance [137]. A large oversight of some *in vitro* biofilms which relate to CF lung infections, is the use of the *P. aeruginosa* reference strain PAO1. The lack of alginate production in PAO1 makes it difficult to gauge the relevance of these studies to CF lung infections. Whilst non-mucoid strains might be present in the CF lung, infection with *P. aeruginosa* is characterized in many instances by the presence of this mucoid phenotype. However, as one of the most studied *P. aeruginosa* strains, its use has been exemplary in identifying and understanding complex regulatory pathways. If we are to use clinical isolates more readily in experiments, these must be chosen with care. Recent studies have shown that in CF infections, although there may be only one infection clone of *P. aeruginosa* (eg: The Liverpool Epidemic Strain, LES), the population of LES can be highly phenotypically diverse [105,138]. More recently, population analysis of the LES, has identified the commonality of divergent sub-lineages and their co-existence, allowing them to exchange potentially adaptive mutations [138]. Put simply, which of these LES isolates would you select to be your choice as ‘the’ LES strain? In this instance, one option is to consider working with ‘populations’ of *P*. *aeruginosa* taken from a sputum sample rather than taking a single colony from a plate. We also wish to highlight the clinical ramifications of such diversity, especially during antimicrobial therapy, whereby sub-inhibitory concentrations can drive diversification which may affect the patients clinical outcome [108]. The extensive use of antibiotic “cocktails” during CF-lung related infections highlights the need to take diversification of populations seriously, and not rely on a single clone or strain. Combining this information, it might be more relevant to utilise clinical isolates from early infections, in a representative media with a clinical antibiotic treatment regime, thus allowing us to study the diversification process and how it impacts various aspects of disease.

If we wish to mimic the *in vivo* environment, then the use of *ex vivo* samples provides us with new opportunities [92,137,139–142]. By using *ex vivo* tissue samples with *in vitro* methods and synthetic media, it may be possible to create controlled environments similar to those seen *in vivo*. Whilst many factors, most importantly an immune response, remain absent, the replacement of an abiotic surface with a biotic one will allow the growth and study of biofilms similar to those seen *in vivo*. Each of these systems which attempt to bridge the knowledge gap between *in vivo* conditions and *in vitro* methods are summarised in Table 3.

**Table 3.**
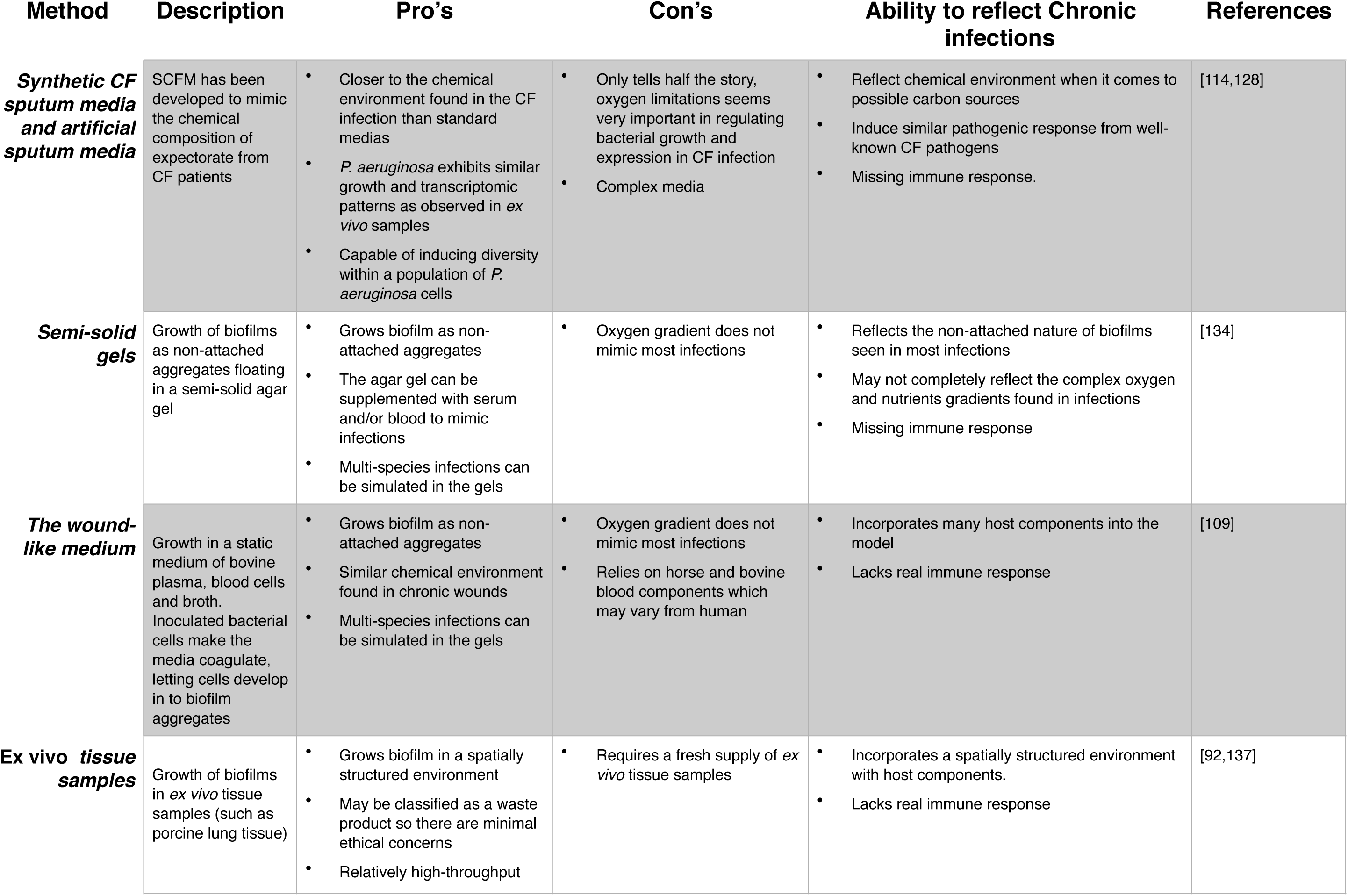
Methods that better represent *in vivo* conditions during *in vitro* investigation

| Method | Description | Pro's | Con's | Ability to reflect Chronic infections | References |
| --- | --- | --- | --- | --- | --- |
| **Synthetic CF sputum media and artificial sputum media** | SCFM has been developed to mimic the chemical composition of expectorate from CF patients | • Closer to the chemical environment found in the CF infection than standard medias<br>• *P. aeruginosa* exhibits similar growth and transcriptomic patterns as observed in *ex vivo* samples<br>• Capable of inducing diversity within a population of *P. aeruginosa* cells | • Only tells half the story, oxygen limitations seems very important in regulating bacterial growth and expression in CF infection<br>• Complex media | • Reflect chemical environment when it comes to possible carbon sources<br>• Induce similar pathogenic response from well-known CF pathogens<br>• Missing immune response. | [114,128] |
| **Semi-solid gels** | Growth of biofilms as non-attached aggregates floating in a semi-solid agar gel | • Grows biofilm as non-attached aggregates<br>• The agar gel can be supplemented with serum and/or blood to mimic infections<br>• Multi-species infections can be simulated in the gels | • Oxygen gradient does not mimic most infections | • Reflects the non-attached nature of biofilms seen in most infections<br>• May not completely reflect the complex oxygen and nutrients gradients found in infections<br>• Missing immune response | [134] |
| **The wound-like medium** | Growth in a static medium of bovine plasma, blood cells and broth. Inoculated bacterial cells make the media coagulate, letting cells develop in to biofilm aggregates | • Grows biofilm as non-attached aggregates<br>• Similar chemical environment found in chronic wounds<br>• Multi-species infections can be simulated in the gels | • Oxygen gradient does not mimic most infections<br>• Relies on horse and bovine blood components which may vary from human | • Incorporates many host components into the model<br>• Lacks real immune response | [109] |
| **Ex vivo tissue samples** | Growth of biofilms in *ex vivo* tissue samples (such as porcine lung tissue) | • Grows biofilm in a spatially structured environment<br>• May be classified as a waste product so there are minimal ethical concerns<br>• Relatively high-throughput | • Requires a fresh supply of *ex vivo* tissue samples | • Incorporates a spatially structured environment with host components.<br>• Lacks real immune response | [92,137] |

## Conclusion and recommendations

To date, most of our mechanistic knowledge and hypotheses surrounding biofilm formation and how this relates to chronic infection is based upon *in vitro* observations, primarily through the use of microtitre plate assays and flow cell systems. These systems have greatly enhanced our knowledge about the mechanisms of how cells attach to surfaces and differentiate into multicellular biofilms. However, it is becoming increasingly apparent that many of our *in vitro* methods do not accurately represent *in vivo* conditions, and so may provide only limited information that has any clinical relevance. Whilst this should come as no surprise, due to the innate complexity of biological systems, it highlights the need for better representation of the *in vivo* condition during *in vitro* investigation. At present, many *in vitro* studies contain (i) unrepresentative nutrients, (ii) uncharacteristic nutrient flow, (iii) uncharacteristic surfaces, and (iv) unrepresentative microorganisms. Whilst there is no “gold-standard” for the study of *in vivo* and *in vitro* biofilm formation, it is crucial to know the limiting factors of selected models so as to not over-extrapolate data, and generate assumptions beyond the capabilities of the model.

It is known that different nutrients alter the way in which bacterial cells grow. The use of synthetic media, which better represents the composition of *in vivo* exudate should be considered to produce data of increased clinical relevance. Other aspects of an *in vivo* micro-environment are more difficult to manipulate, such as the creation of oxygen gradients and a host immune response.

As a research community, we have examined mono-species cultures extensively, systematically, yet not exhaustively, and whilst a leap into polymicrobial research might seem counter intuitive (when multiple aspects of mono-culture are not fully understood), it presents itself as the next logical step. With the majority of chronic infections harboring polymicrobial communities, interactions between different species may critically influence various factors associated with chronic infection such as virulence and AMR. Another issue that we raised is the use of reference strains. Such strains present researchers with a well-defined system which creates some reproducibility of research between laboratories. However, the relevance of these strains to isolates taken directly from infection is often unclear and we should consider whether there is always a clinical relevance to work performed using reference strains. We also know that clinical isolates taken from the same patient can show considerable phenotypic diversity and so there may be a need to use ‘populations’ of bacteria taken from an infection to more accurately study biofilms and infection.

Rome was not built in a day. We cannot expect immediate paradigm shifts in the way experiments are performed, nor do we know how to either. Precise experiments using well defined media and reference strains remain of considerable importance in elucidating mechanisms which might be important for biofilm formation and virulence during infection. The use of representative species, in conjunction with other representative species (creating a poly-microbial “infection”), in media that is representative (such as synthetic sputum media), with representative surfaces (such as an *ex vivo* tissue sample), will likely produce data that is more relevant to *in vivo* infections. Finally, as biologists, we should not be afraid of performing, and being more accepting of, ‘dirty’ experiments. These are experiments where we have less control and stringency, and where some aspects are not 100% standardised. We suggest that this would be a positive way forward in helping us to understand the biology of infection better.

## Acknowledgements

This work was funded by a Human Frontier Science grant to SPD and TB (RGY0081/2012).

